# Maximum entropy models elucidate the contribution of metabolic traits to patterns of community assembly

**DOI:** 10.1101/526764

**Authors:** Jason Bertram, Erica A. Newman, Roderick C. Dewar

## Abstract

**Aim:** Maximum entropy (MaxEnt) models promise a novel approach for understanding community assembly and species abundance patterns. One of these models, the “Maximum Entropy Theory of Ecology” (METE) reproduces many observed species abundance patterns, but is based on an aggregated representation of community structure that does not resolve species identity or explicitly represent species-specific functional traits. In this paper, METE is compared to “Very Entropic Growth” (VEG), a MaxEnt model with a less aggregated representation of community structure that represents species (more correctly, functional types) in terms of their per capita metabolic rates. We examine the contribution of metabolic traits to the patterns of community assembly predicted by VEG and, through aggregation, compare the results with METE predictions in order to gain insight into the biological factors underlying observed patterns of community assembly.

**Innovation:** We formally compare two MaxEnt-based community models, METE and VEG, that differ as to whether or not they represent species-specific functional traits. We empirically test and compare the metabolic predictions of both models, thereby elucidating the role of metabolic traits in patterns of community assembly.

**Main Conclusions:** Our analysis reveals that a key determinant of community metabolic patterns is the “density of species” distribution *ρ*(*ϵ*), where *ρ*(*ϵ*)*dϵ* is the intrinsic number of species with metabolic rates in the range (*ϵ*, *ϵ* + *dϵ*) that are available to a community prior to filtering by environmental constraints. Our analysis suggests that appropriate choice of *ρ*(*ϵ*) in VEG may lead to more realistic predictions than METE, for which *ρ*(*ϵ*) is not defined, and thus opens up new ways to understanding the link between functional traits and patterns of community assembly.

**BIOSKETCHES:** **Jason Bertram** is a mathematical biologist who develops ecological and evolutionary theory at Indiana University.

**Erica A. Newman** is postdoctoral researcher focusing on connections between disturbance ecology and biodiversity patterns, including applied topics in wildlife ecology and wildfire management.

**Roderick C. Dewar** is a professor at the Australian National University, researching entropy-based principles of organization in biological and physical systems.

## 1 INTRODUCTION

One of the central aims of ecology is to understand the determinants of community assembly. Many studies of community assembly involve summaries of community structure such as the species abundance distribution (SAD), species-area relationship (SAR), and analogous metabolic-rate distributions. We will refer to these summary distributions collectively as community structure distributions (CSDs). CSDs, particularly the SAD, have attracted a lot of attention because their shapes are strikingly similar across different communities, representing a rare example of “universality” in community ecology (McGill *et al.* 2007).

The existence of universal features in CSDs is intriguing because these could reflect universal aspects of the biological processes responsible for structuring communities. However, CSDs could also be universal for statistical reasons (Tokeshi 1993; Ulrich *et al*., 2010). Similar to how the normal distribution is ubiquitous because many measured quantities involve statistical averaging (the central limit theorem), CSDs could be universal simply because community-specific details disappear in aggregating patterns to the level of species counts, or other forms of averaging. This would make CSDs considerably less valuable for understanding the biological determinants of community assembly, such as how community structure depends on the functional traits of the organisms in the community (McGill *et al.*, 2006; Díaz et al. 2013). It is therefore important to disentangle the contributions of biological versus statistical factors to CSDs. This issue is closely related to the long-running debate on the relative roles of “mechanism” and “drift” in ecology (McGill and Nekola, 2010; Vellend, 2010), and on ecosystem stability and the role of disturbance (Newman et al. 2018).

A promising recent approach for disentangling biological from statistical factors in ecological models is to use the statistical principle of maximum entropy (MaxEnt). MaxEnt models are “top-down” in that they seek to identify a minimal set of biological assumptions required to reproduce a given empirical pattern (such as a CSD). Once these assumptions have been specified, the MaxEnt principle predicts statistical patterns of community structure by effectively treating all other mechanistic details statistically as unbiased random noise. By empirically testing predictions based on different assumptions, MaxEnt provides a means to resolve the partitioning of biological versus statistical factors in driving observed ecological patterns.

MaxEnt models have had some success at predicting CSDs, but the ecological interpretation of these successes has not been straightforward. A number of MaxEnt models have appeared in the ecological literature with a variety of different assumptions and justifications (*e.g.* Shipley, Vile and Garnier, 2006; Pueyo, He and Zillio, 2007; Harte, Zillio, Conlisk and Smith, 2008; Dewar and Porté, 2008; Banavar, Maritan and Volkov, 2010; Bertram and Dewar, 2015). This had led to extensive debates about the prospects and pitfalls of this approach.

Two key issues in these debates may be identified: one conceptual, the other more technical. The conceptual issue concerns the interpretation of the MaxEnt procedure itself, including the challenge of connecting MaxEnt to familiar ecological processes such as dispersal, disturbance, and interactions between organisms. This issue has been discussed at length elsewhere (Dewar, 2009; McGill and Nekola, 2010; Shipley, 2010; Supp, Xiao, Ernest and White, 2012; Harte and Newman, 2014; Supp and Ernest 2014; Bertram and Dewar, 2015; Newman et al., 2018), and will thus not be our focus here.

Rather, our focus will be on the technical issue, which concerns the level of detail at which the community is described in the model before MaxEnt is even applied (He, 2010; Favretti, 2017). Changing the variables used to describe a community (*e.g.* resolving a community in greater detail) can dramatically alter the predictions that MaxEnt makes about the community, and yet there is no apparent *a priori* reason to prefer one choice of variables over another. As a result, a variety of choices have appeared in different models, usually with little justification. A comparison of these disparate approaches is required in order to better guide the application of MaxEnt models in ecology.

Here we present a comparison of two MaxEnt-based models in ecology which have both successfully reproduced observed CSDs: METE (Maximum Entropy Theory of Ecology; Harte et al. 2008; Harte et al. 2009; Harte 2011; Harte and Newman, 2014) and VEG (Very Entropic Growth; Dewar and Porté, 2008; Bertram and Dewar, 2013; Bertram and Dewar 2015). These models are well suited for our objective of comparison because METE describes communities at the same coarse-grained level of detail as the SAD, whereas VEG is more detailed in that it resolves the abundance of each separate species.

Crucially, this difference in community description allow us to explore the biological determinants of patterns of community assembly. Specifically, VEG distinguishes species by their per capita metabolic traits. By contrast, METE only distinguishes separate species by their abundances, and requires the total number of species present in the community as an input rather than as a prediction. METE then predicts a distribution of metabolic rates for the individuals in a species as a function of its abundance, imparting functional traits statistically by abundance, rather than by species identity (Section 2.1). It is therefore not possible to investigate how different species-specific functional traits might modify community structure using METE. Thus, by comparing METE and VEG, we are able to investigate more transparently what sorts of functional trait assumptions are necessary for reproducing observed patterns.

## 2 METE AND VEG: TWO MAXENT MODELS OF COMMUNITY ASSEMBLY

### 2.1 METE

The central quantity predicted by METE is a joint probability distribution *R*_*M*_(*n*, *ϵ*) called the “ecosystem structure function.” By definition, *R*_*M*_(*n*, *ϵ*)*dϵ* is the joint probability that a species selected at random from a community has abundance *n*, and that an individual selected at random from a species with abundance *n* has a metabolic requirement between *ϵ* and *ϵ* + *dϵ* (Harte et al. 2008; Harte, 2011; Appendix A). The ecosystem structure function is closely related to the SAD: if we add together all of the possible metabolic requirements *ϵ* we obtain the probability distribution for the abundance of a randomly selected species, *R*_*M*_(*n*) = ∫ *R*_*M*_(*n*, *ϵ*)*dϵ*; the SAD is then simply *SR*_*M*_(*n*) where *S* is the total number of species present in the community. Thus, *R*_*M*_(*n*, *ϵ*) is a SAD that has been extended to also incorporate information about community metabolic structure.

METE assumes that *R*_*M*_(*n*, *ϵ*) satisfies two constraints

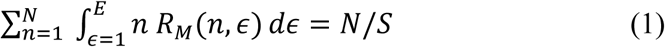

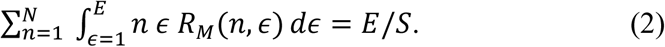

In words, these constraints say that the total number of individuals in the community 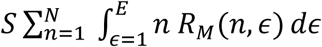 is equal to *N*, and the total community metabolic requirement 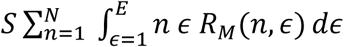 is equal to *E*. *R*_*M*_(*n*, *ϵ*) is then obtained by maximizing the Shannon entropy −∑_*n*_ ∫ *R*_*M*_ ln *R*_*M*_ ln *R*_*M dϵ*_ subject to constraints (1) and (2), as well the constraint that *R*_*M*_(*n*, *ϵ*) sums to 1 (since it is a probability distribution). This maximization procedure gives

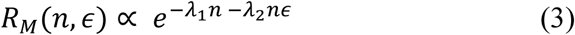

where *𝜆*_1_ and *𝜆*_2_ are constants (Lagrange multipliers) with values chosen such that constraints (1) and (2) hold (for details, see Harte, 2011). The triplet of values *N*, *E* and *S* are the inputs to METE (note that we will not consider the area-scaling component of the full METE theory; Harte, 2011).

### 2.2 VEG

VEG is similar to METE in that it uses MaxEnt to infer community properties from a few constraints. Moreover, the VEG constraints are similar to METE’s (see below). The major feature that differentiates VEG is that it represents community structure in more detail. In VEG, species are distinguishable, whereas METE only specifies the proportion of species with each abundance *n* via the ecosystem structure function (Fig. 1).

**Figure 1.**
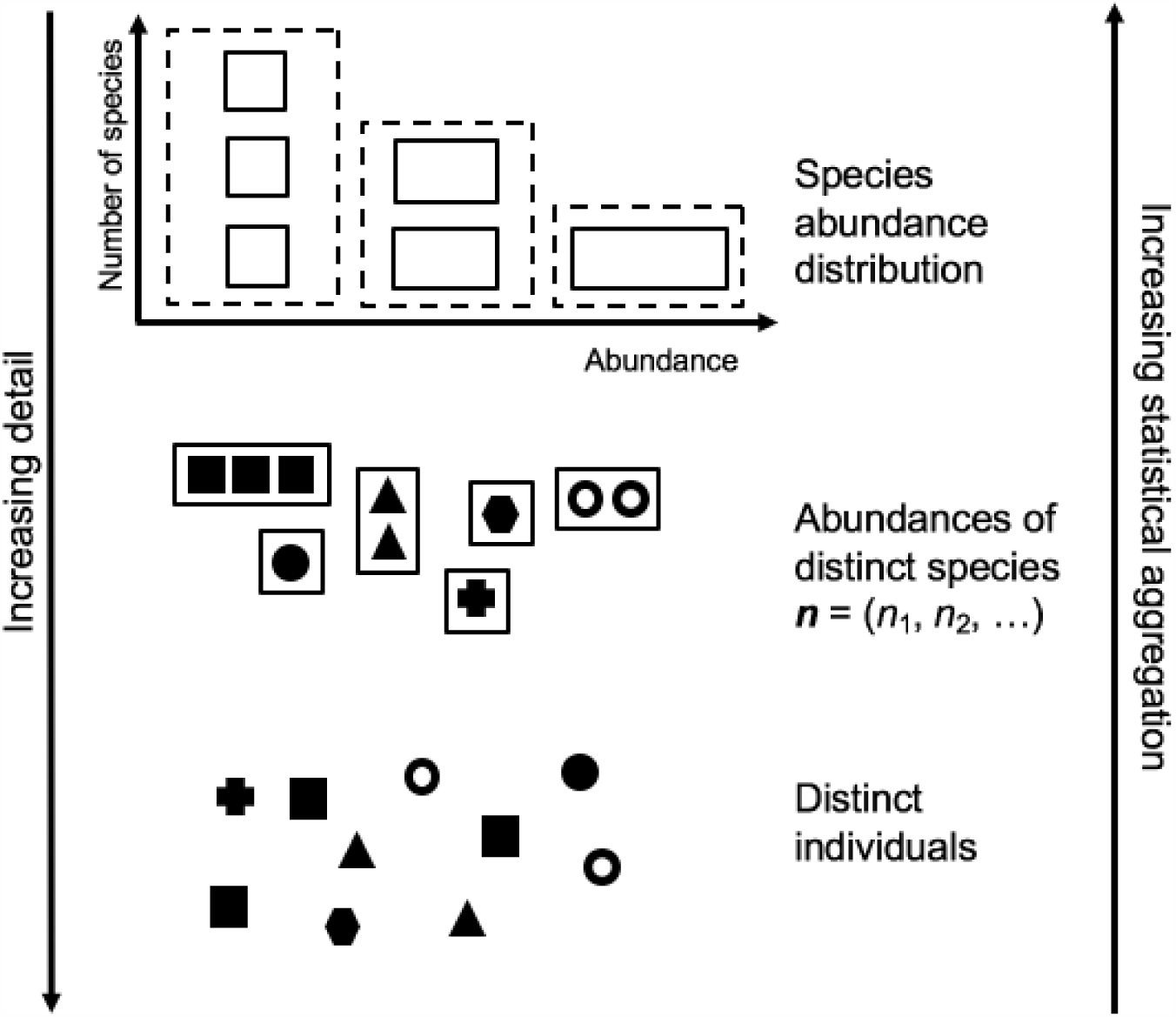
Three levels of detail commonly used for describing ecological communities. At the greatest level of detail (bottom), the distinct identities of individuals and their spatial locations are known. At the intermediate levels of detail found in many well-mixed models such as Lotka-Volterra models (middle), the abundance of each distinct species is known. At the lowest level of detail and highest level of statistical aggregation (top), species identities are lost, and the SAD provides the only description of species diversity.

In contrast to METE (which uses MaxEnt to infer the ecosystem structure function directly), VEG uses MaxEnt to predict the probability *p*(***n***) that, when we take a snapshot of the community, we observe the species abundances ***n*** = (*n*_1_, *n*_2_, …) (*i.e.* the species labeled 1 has abundance *n*_1_, and so on). In VEG, species’ abundances may be zero; the number of species actually present in a snapshot is the number of nonzero elements of ***n***. VEG therefore predicts probabilities for the abundance of each species separately; consequently, VEG also predicts the expected number species that are present in the community. Species in VEG are also assigned distinct functional traits: the individuals of species *i*are assumed to have a metabolic requirement of *ϵ*_*i*_, where the species labels are chosen such that *ϵ*_1_ ≤ *ϵ*_2_ ≤ *ϵ*_3_ …, and so on.

Similar to METE, VEG assumes total abundance and total metabolic requirement constraints

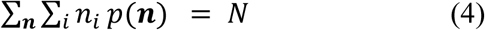

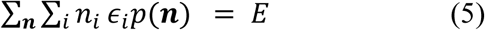

Since *p*(***n***) represents the probability of observing the “snapshots” ***n***, the probabilities can be interpreted as sample frequencies representing the proportion of time that the community spends with different abundance compositions ***n***. Consequently, constraints (4) and (5) have a clear ecological interpretation in VEG as fixing the time-averaged total abundance and total metabolic requirement of the community to have the values *N* and *E* respectively; the latter can be interpreted as an expression of the long-term steady-state ecological balance between resource use (left-hand side of Eq. (5)) and supply (*E*, right-hand side of Eq. (5)). In contrast, the METE constraints (Eqs. (1) and (2)), which are statements about “information”, do not have a similarly straightforward ecological interpretation.

Again similarly to METE, *p*(***n***) is obtained by maximizing the Shannon entropy **−** ∑_***n***_ *p*(***n***) ln *p*(***n***) subject to constraints (4), (5), and the constraint that *p*(***n***) sums to 1. This maximization procedure gives

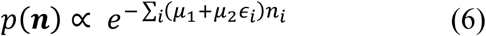

where *μ*_1_ and *μ*_2_ are the Lagrange multipliers corresponding to constraints (4) and (5) respectively. Note that in Eq. (6), *p*(***n***) depends on the spectrum of metabolic requirements present in the community *ϵ*_1_ ≤ *ϵ*_2_ ≤ ⋯. Thus the inputs of VEG are *N*, *E* and the spectrum of values *ϵ*_*i*_. In contrast to METE, the number of species (*S*) present in the community is an output of VEG, rather than an input.

## 3 COMPARING METE AND VEG

### 3.1 THE VEG ECOSYSTEM STRUCTURE FUNCTION

In this section we give an intuitive derivation of the ecosystem structure function implied by VEG, which will be denoted *R*_*V*_(*n*, *ϵ*) (a more rigorous mathematical derivation is given in Appendix B). This will allow us to directly compare the predictions of METE and VEG.

When we sample a species at random from a community, all species present have the same probability of being selected. However, the metabolic requirement *ϵ* of the selected species is more likely to take some values than others due to two effects: (1) *Trait availability.* Among the species currently inhabiting the community’s broader geographic region, some values of *ϵ* are more likely to occur than others due to intrinsic biophysical constraints on the traits determining *ϵ*, and the region’s evolutionary history; (2) *Environmental filtering* (Shipley *et al.*, 2006). From the distribution of possible metabolic rates, some values of *ϵ* are more likely to be actually present in the community due to additional bias imposed by local environmental constraints (such as Eqs. (4) and (5)).

VEG represents a special case in which there is no trait variation within species: all individuals in species *i*have the same metabolic requirement *ϵ*_*i*_ (thus a VEG “species” is more appropriately interpreted as a functional type rather than a taxonomic unit; Bertram and Dewar, 2013). Thus, the first effect above (trait availability) is represented by the fact that the metabolic spectrum *ϵ*_1_ ≤ *ϵ*_2_ ≤ ⋯ may be more densely packed at some values of *ϵ* than at others. To represent this effect mathematically, we introduce the “density of species” distribution *ρ*(*ϵ*); *ρ*(*ϵ*)*dϵ* counts the number of metabolic requirement values (“species”) contained in the interval (*ϵ*, *ϵ* + *dϵ*). For comparison with METE, in which *ϵ* is a continuous variable, we assume that the metabolic requirement spectrum is sufficiently dense that we can approximate *ρ*(*ϵ*) as a continuous function of *ϵ*. Intuitively, the shape of *ρ*(*ϵ*) represents the relative probabilities that a species selected at random out of all possible species that could be present in the community has a metabolic requirement within a given interval (Fig. 2).

**Figure 2.**
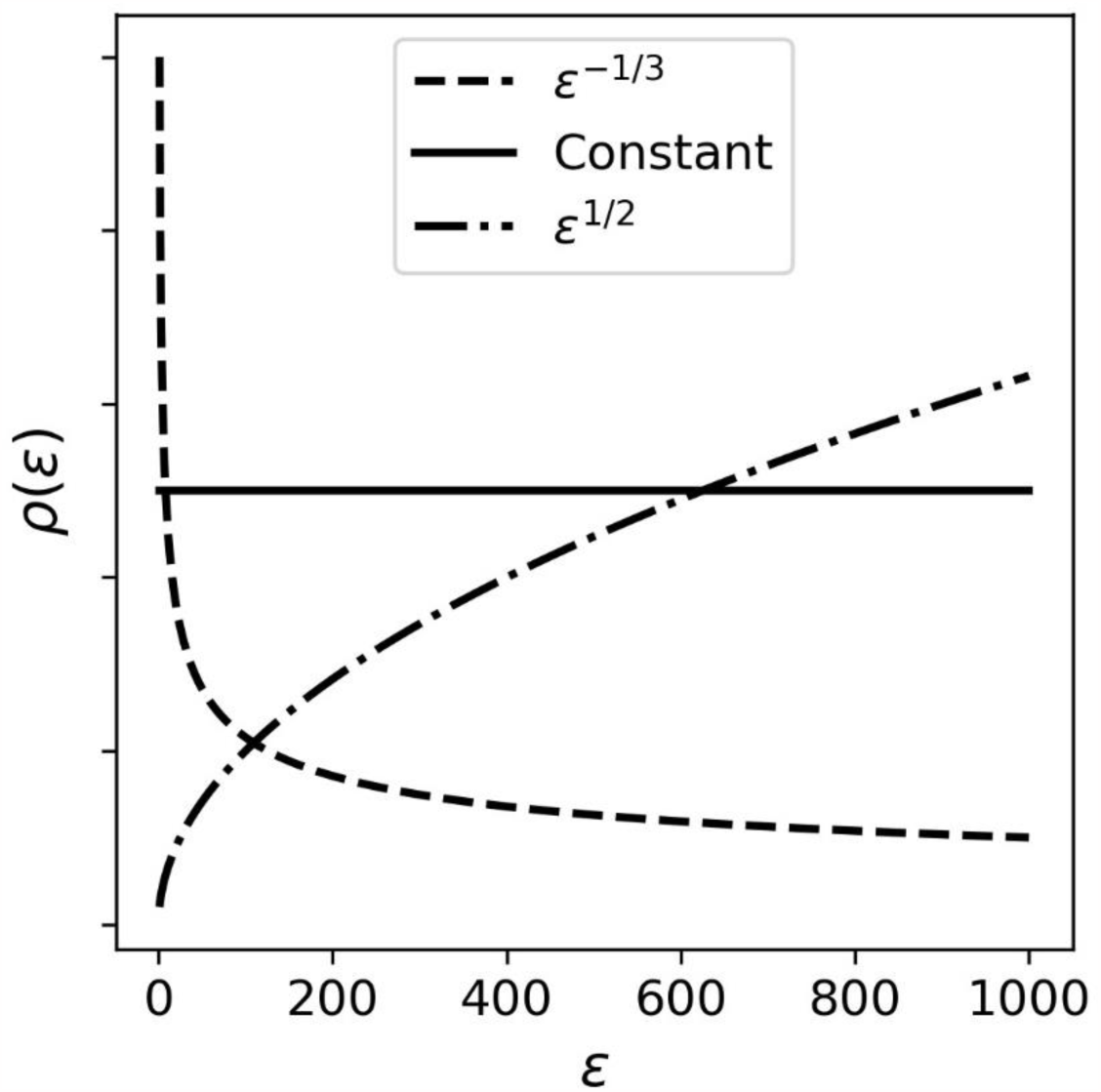
The function *ρ*(*ϵ*) counts the local density of metabolic rates in the assumed spectrum of possible rates *ϵ*_1_ ≤ *ϵ*_2_ ≤ ⋯ in VEG. It represents the relatively probability that a randomly selected VEG “species” has metabolic rate *ϵ* when sampled from all possible “species”.

Once a species has been sampled out of all possible species and its metabolic requirement has been found to be *ϵ*, the probability that it has abundance *n*, denoted *p*(*n*|*ϵ*), can then be straightforwardly calculated in VEG from Eq. (6) (from Appendix B, *p*(*n*|*ϵ*) ∝ *e* ^−(*μ*1+*μ*2*ϵ*)*n*^). VEG also explicitly accounts for the second effect above (environmental filtering), through the Lagrange multipliers *μ*_1_ and *μ*_2_ that reflect the environmental constraints of Eqs. (4) and (5).

To construct *R*_*V*_(*n*, *ϵ*), which only refers to species that are actually present, we restrict our attention to *n* ≥ 1. Thus, the joint probability of sampling a species with abundance *n* from the community, and an individual from such a species with metabolic requirement *ϵ*, is proportional to *ρ*(*ϵ*)*p*(*n*|*ϵ*), where *n* ≥1. This gives

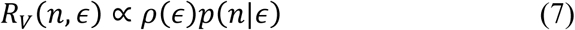

The above argument leading to Eq. (7) for *R*_*V*_ (and the more rigorous argument given in Appendix B) is quite general. It can be applied to obtain the ecosystem structure function for any model in which we know the density of species *ρ*(*ϵ*) (which need not be restricted to a species-specific trait spectrum as in VEG), and which predicts abundance probabilities conditional on the trait values *p*(*n*|*ϵ*) (whether those probabilities are predicted using MaxEnt or by other means).

### 3.2 SPECIES ABUNDANCE DISTRIBUTIONS

A large number of ecological models have reproduced realistic SADs, including METE (Harte et al., 2008) and VEG (Bertram and Dewar, 2015). SAD comparisons consequently only have weak power to discriminate the predictions of different ecological theories (they are “weak tests”; McGill 2003, McGill et al. 2007). In particular, the SAD predictions of METE and VEG will not tell us much about their differences. It is interesting to demonstrate this “weak test” property of SADs explicitly in terms of the METE and VEG ecosystem structure functions.

As noted in section 2.1, the SAD is obtained by integrating the ecosystem structure function over *ϵ* (the SAD is proportional to *R*(*n*) = ∫ *R*(*n*, *ϵ*)*dϵ*). We therefore expect that the SAD will be to some extent insensitive to the exact manner in which *R*(*n*, *ϵ*) depends on *ϵ*.

In the case of METE, the predicted SAD is almost entirely independent of the value of *E*/*S* in the metabolic constraint Eq. (2) for many of the most heavily studied SAD datasets (i.e. *R*_*M*_(*n*) is independent of *λ*_2_; Harte et al. 2008). This behavior represents the limiting case of large *E*, corresponding to resource-rich communities. Thus, in many cases of interest, METE produces SADs that are insensitive to the value of *E*/*S* (note, however, that the existence of the metabolic constraint Eq. (2) is necessary to get a Fisher log-series form for 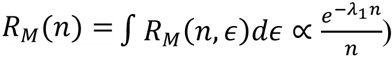.

VEG allows us investigate the “weak test” property in greater depth because we can independently change the form of *ρ*(*ϵ*) and check if this appreciably changes the VEG SAD. Suppose for illustrative purposes that the metabolic requirement spectrum has a power law form (Dewar and Porté, 2008)

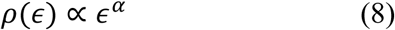

where *α* is a free parameter. Our motivation for Eq. (8) is to have a simple one-parameter function in which we can control the relative density of species at low versus high *ϵ* (*α* = 0 corresponds to a uniformly spaced spectrum; Fig. 2). Using Eq. (8), it can be shown that

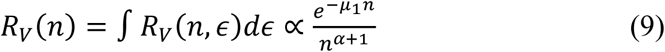

for all but the lowest abundance species (see Appendix C). Thus, although the exact quantitative shape of *R*_*V*_(*n*) does depend on *α* (both explicitly in Eq. (9) and implicitly via the fact that *μ*_1_ and *μ*_2_ depend on *α*), *R*_*V*_(*n*) will qualitatively have the familiar “hollow curve” shape (McGill et al. 2007) regardless of the particular choice of *α*. In particular, *α* = 0 gives the Fisher log-series (similar to METE). Thus, since *ρ*(*ϵ*) represents the spectrum of functional traits, we can conclude that the shape of the VEG SAD is only marginally sensitive to the metabolic trait values of the species present.

However, recall that VEG predicts the total number of species/functional types *S* in the community (Sec. 2.2). This predicted *S* is more sensitive to the assumed metabolic trait values than the SAD shape, and could differ from the observed value of *S* for given observed values of *N* and *E*. By contrast, METE uses the empirically observed value of *S* to construct the METE SAD.

### 3.3 METABOLIC-RANK DISTRIBUTIONS

In this section we compare the metabolic dependence of the two structure functions *R*_*V*_(*n*, *ϵ*) and *R*_*M*_(*n*, *ϵ*). We do this in two ways: via the marginal distribution for individual metabolic rates *R*(*ϵ*) = ∑_*n*_*R*(*n*, *ϵ*), and via the individual-level energy distribution (IED) defined by 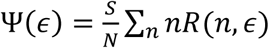 (Harte 2011; Newman *et al.*, 2014). *R*(*ϵ*) is the probability that a species sampled at random from the community has metabolic rate *ϵ*, while Ψ(*ϵ*) is the probability that an individual sampled at random from the community has metabolic requirement *ϵ*. In contrast to the SAD (Section 3.1), *R*(*ϵ*) and Ψ(*ϵ*) are both sensitive to the shape of *ρ*(*ϵ*) in VEG. We can thus ask, what shape does *ρ*(*ϵ*) need to be to match metabolic data, and how does this *ρ*(*ϵ*) compare to the predictions of METE?

Following Harte et al. (2017), we calculate and plot Ψ(*ϵ*) cumulatively such that log metabolic rate appears on the vertical axis, and the horizontal axis is the proportion of the population with metabolic rate greater than or equal to a given *ϵ* (*i.e.* the rank of the corresponding individual). We assumed a power law spectral density as in Eq. (8), taking *α* as a free parameter to be fitted, and then minimized the least-squares difference between measured log metabolic rates and the predictions of VEG. We repeated this procedure for the three datasets considered in Harte et al. (2017): Barro Colorado Island trees (Hubbell et al. 2005), Hawaiian island arthropods (Gruner, 2007), and Rocky Mountain subalpine meadow plants (Newman et al. 2014).

In all three datasets we found values of *α* that give superior Ψ(*ϵ*) fits to METE (bottom three panels of Fig. 3; note the logarithmic horizontal axis). This is no great victory given that we have introduced a free parameter *α* that is not available to METE, but it confirms that the power law form for *ρ*(*ϵ*) gives plausible metabolic predictions. METE and VEG both track the middle and higher ranks closely, but at lower ranks the VEG metabolic rates are too low whereas the METE predictions are too high. The corresponding marginal metabolic distributions *R*_*M*_(*ϵ*) and *R*_*V*_(*ϵ*) (upper panels in Fig. 3) confirm that METE assigns higher probabilities to the highest values of *ϵ* (*R*_*M*_(*ϵ*) has a longer tail).

**Figure 3.**
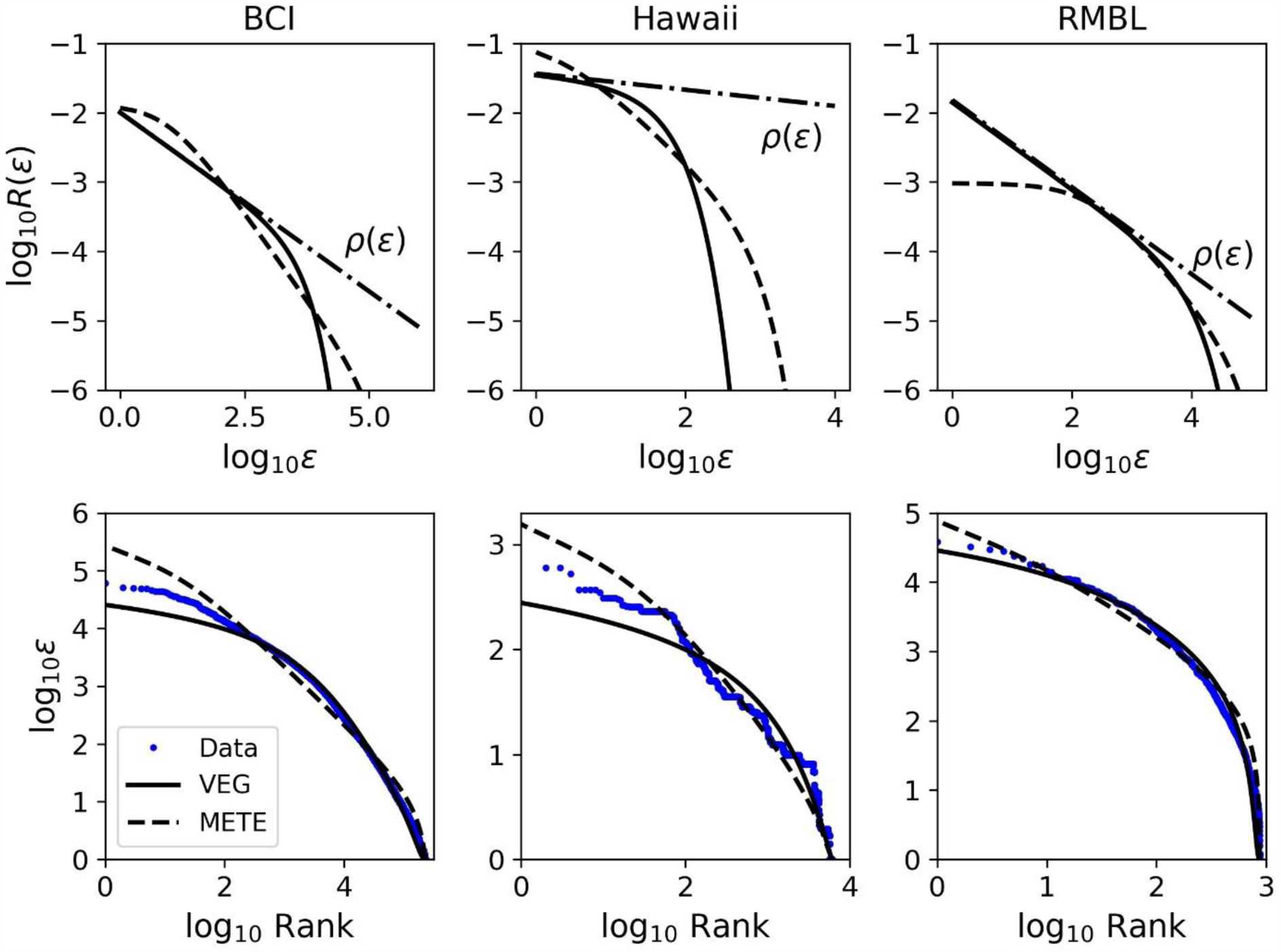
Comparison of METE and VEG rank-metabolism relationships in multiple communities: Barro Colorado Island trees (BCI; *α* = −0.52, *ρ*(1) = 10^4^), Hawaiian island arthropods (Hawaii; *α* = −0.12, *ρ*(1) = 10^2^), and Rocky Mountain subalpine meadow plants (RMBL; *α* = −0.63, *ρ*(1) = 10^2^) communities. Each upper/lower panel pair shows the *R*(*ϵ*) and rank-*ϵ* curves for the same METE and VEG ecosystem structure functions.

## 4 DISCUSSION

A key insight of the above analysis is that the ecosystem structure *R*(*n*, *ϵ*) is sensitive to the shape of the density of species distribution represented mathematically by *ρ*(*ϵ*). In the case of VEG, Eq. (7) implies *R*_*V*_(*ϵ*) ∝ *ρ*(*ϵ*) ∑_*n*≥1_ *p*(*n*|*ϵ*) = *ρ*(*ϵ*)[1 − *p*(0|*ϵ*)]. This expression clearly shows the two effects introduced at the start of Sec. 3.1: *R*(*ϵ*) is the density of species *ρ*(*ϵ*) multiplied by the probability 1 − *p*(0|*ϵ*) that a species with metabolic rate *ϵ* is actually present in the community.

Whereas VEG requires us to specify the form of *ρ*(*ϵ*), METE infers *R*_*M*_(*n*, *ϵ*) using only MaxEnt and the constraint equations (1) and (2). In this sense METE is a null model for the contribution of functional traits to community patterns, treating functional traits as “random noise” within the community constraints imposed by *S*, *N*, and *E*. However, METE only infers the trait distribution as would be observed in already-assembled communities. METE refers only to species that are already present in the community, and does not give an expression for *p*(0|*ϵ*); it is therefore not possible to compute the density of species *ρ*(*ϵ*) implicitly inferred by METE. Nonetheless, observed ecological communities generally have a large proportion of individuals with low metabolic requirement. This implies *p*(0|*ϵ*) ≈ 0 and thus *R*(*ϵ*) ≈ *ρ*(*ϵ*) for low *ϵ* (see the convergence of *R*(*ϵ*) and *ρ*(*ϵ*) in VEG in the upper panels of Fig. 3), giving us a glimpse of the *ρ*(*ϵ*) predictions of METE.

VEG explicitly separates the trait values that are possible from the trait values that are actually observed post-assembly. Since *ρ*(*ϵ*) is an input, VEG represents an explicit model for the contribution of functional traits to CSDs. This begs the question of what then determines *ρ*(*ϵ*) as the appropriate choice in VEG. There are at least two answers:

(i) On short timescales, *ρ*(*ϵ*) may simply express the mix of potential species that are available to the community at any given time, as in biodiversity manipulation experiments where a given restricted set of species is thrown together and left to self-organize. This short-term *ρ*(*ϵ*) could be highly contingent on the community’s recent history, and could have a strong effect on metabolic patterns following disturbance.
(ii) On longer timescales, *ρ*(*ϵ*) may express the totality of conceivable species that might be available to the community. In this case *ρ*(*ϵ*) would depend on how we define species in the first place. With reference to Eq. (7), the choice *α* = 0 corresponds to *defining* “species” by their metabolic requirement, *i.e.* discretize *ϵ*–space into equal intervals of width ∆*ϵ* and define species *i* to be the set of individuals whose metabolic requirement *e* lies between (*i*− 1)∆*e* and *i*∆*e*. Alternatively, “species” could be defined via biomass (in which case the value of *α* in Eq. (8) may reflect metabolic scaling as in Dewar & Porté 2008); or via other individual traits (*t*) on which metabolic requirement depends, *ϵ*(*t*).

In either case, VEG opens up ways to understanding the link between functional traits and CSDs that are simply not available to METE.

One of METE’s great strengths is that it only requires three parameters *S*, *N* and *E* for all of its predictions. How might the above insights be used to improve the predictions of METE without damaging this exceptional parsimony? The answer may lie in the inclusion of a prior distribution for *ϵ* representing a contribution from the density of species *ρ*(*ϵ*), which plays a role in ecology analogous to the “density of states” in physics describing the distribution in energy-space of available quantum-mechanical particle states. The upper panels of Fig. 3 suggest that this trait distribution should give less weight to the higher values of *ϵ*, and also to the lower values of *ϵ* in the Barro Colorado and Hawaiian communities.

## APPENDIX A: NOTE ON THE DEFINITION OF THE ECOSYSTEM STRUCTURE FUNCTION

Our definition of the ecosystem structure function differs slightly from that given in Harte et al. (2008), which reads “[The ecosystem structure function] is the probability that if a species is picked at random […], then it has abundance *n* and if an individual is picked at random *from that species*, then its metabolic requirement is in the range *ϵ*, *ϵ* + *dϵ*” (our italics). The Harte et al. (2008) definition suggests that METE keeps track of species identity. In fact, *R*_*M*_(*n*, *ϵ*) depends only on *n* and *ϵ*, and not species identity. Thus, *R*_*M*_(*ϵ*|*n*) = *R*_*M*_(*n*, *ϵ*)/ ∑_*n*_ *R*_*M*_(*n*, *ϵ*) is the probability of picking an individual with metabolic requirement *ϵ* conditional on it coming from a species with abundance *n*. There is no way within METE to distinguish between different species with the same abundance *n*, and therefore there is no reason to specify which species the individual is selected from in the definition of the ecosystem structure function (Favretti, 2017).

## APPENDIX B: DERIVING THE ECOSYSTEM STRUCTURE FUNCTION FROM DISTINGUISHABLE SPECIES

In section 2.1, the METE structure function *R*_*M*_ was inferred directly from community-level constraints. Here we derive an analogous VEG structure function *R*_*V*_(*n*, *ϵ*).

We start by defining the probability distribution *P*(*n*, *ϵ*, *i*, ***n***) as follows: *P*(*n*, *ϵ*, *i*, ***n***)*dϵ* is the joint probability that the community has species abundances ***n***, that a species picked at random from the community has species label *i*, that this chosen species has abundance *n*_*i*_ = *n*, and that an individual from this chosen species has metabolic requirement in the interval (*ϵ*, *ϵ* + *dϵ*). *R*_*V*_(*n*, *ϵ*) is then obtained by marginalizing with respect to *i*and ***n***, i.e. *R*_*V*_(*n*, *ϵ*) = ∑_*i*_∑_***n***_ *P*(*n*, *ϵ*, *i*, ***n***) where *n* ≥ 1.

To marginalize *P*, we first write it as a product of conditional distributions

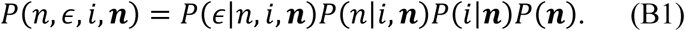

Here *P*(***n***) = *p*(***n***) is the probability that the community has abundance vector ***n***, 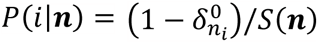 is the probability that a species picked from a community with abundances ***n*** has species label *i*(i.e. 0 if species *i*is absent, 1/*S*(***n***) if present), 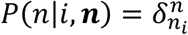 is the probabilty that species *i*has abundance *n* given the species abundances are ***n***, and *P*(*ϵ*|*n*, *i*, ***n***)*dϵ* is the probability that an individual picked from species *i*has metabolic requirement in the interval (*ϵ*, *ϵ* + *dϵ*) given species *i*has abundance *n* and the community abundances are ***n***. We thus obtain

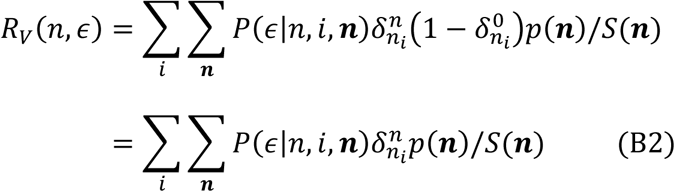

where we have used the fact that 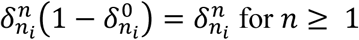 for *n* ≥ 1.

In the case of VEG, all individuals in species *i* have the same metabolic requirement *ϵ*_*i*_, and so *P*(*ϵ*|*n*, *i*, ***n***) = *δ*(*ϵ* − *ϵ*_*i*_) where *δ* is the Dirac delta function (i.e. the probability that a randomly selected individual from species *i*has metabolic requirement *ϵ* is 1 in the immediate vicinity of *ϵ*_*i*_, and is 0 otherwise). Thus, from (B2) we have

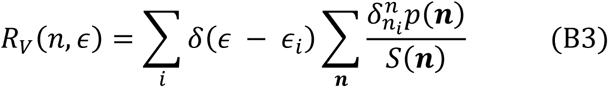

An ecologically important special case of (B3) occurs when the variation in the number of species present from one snapshot to the next is small relative to the expected number of species, such that *S*(***n***) is approximately constant with value given by *S* = ∑_***n***_ *S*(***n***) *p*(***n***). This occurs, for example, if most of the species present have large expected abundances. Eq. (B3) then simplifies to

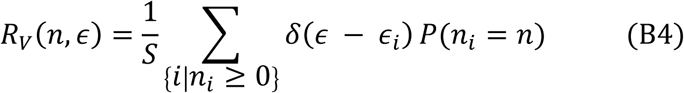

where 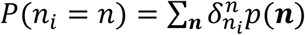 is the probability that species *i*has abundance *n*.

In VEG, we have from Eq. (6) (Bertram and Dewar, 2015)

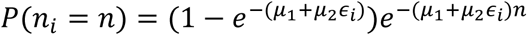

To make it explicit that this probability depends on the metabolic requirement of species *i*, we use the notation *p*(*n*|*ϵ*_*i*_) ≡ *P*(*n*_*i*_ = *n*) (that is, *P*(*n*_*i*_ = *n*) in VEG is the probability that a species has abundance *n* given that its metabolic requirement is *ϵ*_*i*_).

Since the ecosystem structure function is a probability density in the continuous variable *ϵ*, we can introduce a spectral density *ρ*(*ϵ*)*dϵ* that counts the number of metabolic requirement levels *ϵ*_*i*_ in each interval (*ϵ*, *ϵ* + *dϵ*). From Eq. (B4), *R*_*V*_(*n*, *ϵ*) can then be written in the form

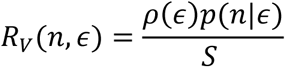

## APPENDIX C: THE VEG SPECIES ABUNDANCE DISTRIBUTION

Assuming a power-law density of states *ρ*(*ϵ*) ∝ *ϵ*^*α*^, we have from Eq. (7)

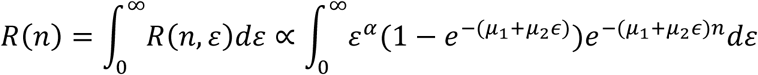

By making the substitution *x* = *μ*_2_*εn*, the integral is found to be:

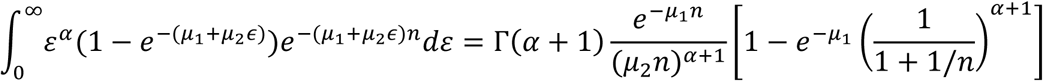

where

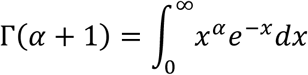

is the gamma function.

For large *n* we have 1/(1 + 1/*n*) ≈ 1 so that:

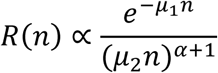

## ACKNOWLEDGEMENTS

Funding for JB was provided by the Environmental Resilience Institute. Postdoctoral funding for EAN was provided by University of Arizona Bridging Biodiversity and Conservation Science program, and the USDA Forest Service. We thank John Harte for comments on earlier versions of this manuscript, and Dan Gruner for use of his dataset.

